# Resolving hidden stoichiometries in Bacterial proteasome activator (Bpa)-substrate complexes by cryo-EM and charge detection mass spectrometry

**DOI:** 10.64898/2026.05.01.722288

**Authors:** Bradley T.V. Davis, Adwaith B. Uday, Anisha Haris, Alexander F. A. Keszei, Jakub Ujma, David Bruton, Kieth Richardson, Mohammad Mazhab-Jafari, Kevin Giles, Natalie Zeytuni, Siavash Vahidi

## Abstract

Bpa (<u>B</u>acterial <u>p</u>roteasome <u>a</u>ctivator) is a regulatory particle within the *Mycobacterium tuberculosis* (*Mtb*) proteasome system that that facilitates ATP-independent substrate engagement and delivery to the 20S core particle (CP) for degradation. The best characterized Bpa substrate is HspR, a transcriptional repressor of *Mtb* stress-response genes whose Bpa-dependent degradation is required for pathogen virulence. However, the stoichiometry of the Bpa:HspR complex, the molecular mechanism of substrate engagement, and the heterogeneity of the resulting assemblies remain unclear. Here, we combine charge detection mass spectrometry (CDMS) and single-particle electron cryomicroscopy (cryo-EM) as complementary approaches to characterize both apo and HspR-bound Bpa. CDMS revealed a previously unreported undecameric apo species and defines a Bpa_12_:HspR_2_ complex stoichiometry with minimal heterogeneity. Cryo-EM, performed without the employment of cross-linking reagents, independently confirmed the presence of undecameric Bpa in solution and localized substrate-associated density to the C-terminal H4 helix of Bpa. Together, these complementary single-particle approaches inform future efforts to target the *Mtb* proteasome system and provide new molecular insight into proteasomal substrate recognition in prokaryotes.

**Table of Content Graphic:** 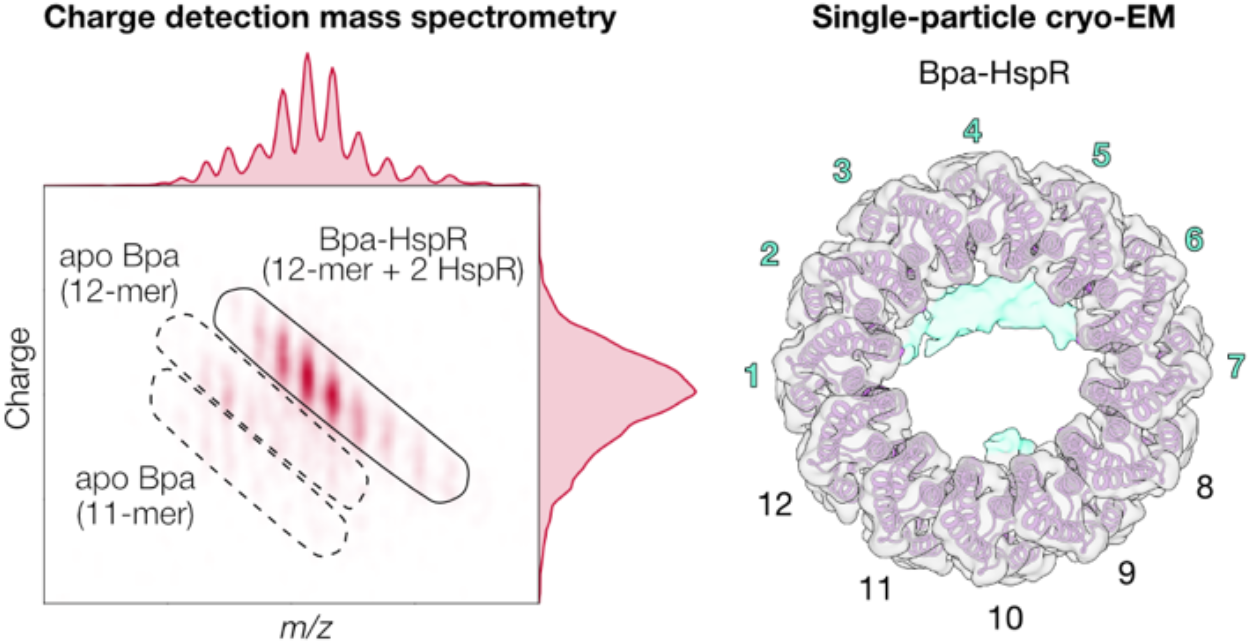

## Introduction

Determining the stoichiometry and heterogeneity of biomolecular assemblies is essential for understanding their function, yet many biologically important complexes resist characterization by conventional structural and biophysical methods. Charge detection mass spectrometry (CDMS) is emerging as a uniquely powerful approach for resolving the stoichiometries of heterogeneous biomolecular assemblies that challenge conventional tools^1^. In electrospray (ESI)-based MS experiments, most macromolecular analytes give rise to a charge state distribution, where a single molecular species with a precisely defined mass appears as a series of peaks along the *m/z* measurement axis, due to different numbers of charges attached to otherwise isobaric molecules^2^. Accurate mass determination therefore relies critically on correct assignment of the underlying charge states^3^. This approach is effective when co-existing molecular species occupy distinct *m/z* regions and when charge state series are well separated^4^. However, many biologically relevant assemblies violate these criteria^5–7^. Closely related oligomers can produce nearly identical *m/z* envelopes, and changes in charge state distributions can mask or mimic changes in mass^8^. As a result, mixtures of stoichiometries may appear deceptively homogeneous, and minor populations can remain experimentally inaccessible^9^. These limitations are especially acute for complexes in the megadalton mass range, where broad charge state distributions, mass heterogeneity, and incomplete desolvation obscure distinct charge states and complicate robust deconvolution of *m/z*-based spectra. Similar challenges also arise for smaller heterogeneous assemblies when closely related stoichiometries or overlapping charge state distributions are present. Orthogonal solution-based methods such as size-exclusion chromatography coupled to multi-angle light scattering (SEC-MALS) and other spectroscopic tools cannot resolve species with similar hydrodynamic radii and light-scattering properties. CDMS addresses this need by independently measuring both *m/z* and charge (*z*) for single ions and computing mass directly from these two measurables^10–15^. Unlike conventional native MS, CDMS does not require resolution of discrete charge states in the *m/z* dimension to obtain information on mass. Instead, by measuring *z* explicitly, CDMS can separate co-existing ion populations that overlap in *m/z* space. This capability is particularly valuable for mixtures in which charge state series overlap, and for systems where low-abundance species are masked by dominant populations.

Here we apply CDMS to the *Mycobacterium tuberculosis* (*Mtb*) proteasome system. The mycobacterial proteasome system is an attractive target for understanding pathogen stress resistance and for exploring new therapeutic strategies^16–20^. The *Mtb* proteasome system is comprised of the 20S core particle (CP) and regulatory particles (RPs) that modulate substrate degradation. The *Mtb* 20S CP adopts a α_7_-β_7_-β_7_-α_7_ configuration that sequesters fourteen catalytic sites within its homoheptameric β-rings. A gating mechanism within the α_7_-rings of the 20S CP restricts substrate access and spurious degradation^21^. Targeted proteolysis requires regulatory particles (RPs) that bind the α_7_-rings via C-terminal GQYL motifs to facilitate gate opening and enable proteasomal degradation of large folded, partially folded, or unfolded substrates^22–27^. Bpa (<u>B</u>acterial <u>p</u>roteasome <u>a</u>ctivator, *Rv3780*) facilitates ATP-independent degradation of a distinct set of substrates^22,25^, most notably HspR, a transcriptional repressor that negatively regulates the expression of the Hsp70/Hsp40 chaperone operon and the associated stress-response genes in *Mtb*^20,25,28,29^. Although HspR is the most extensively characterized Bpa substrate, no experimental structures of apo or bound HspR is available, leaving the molecular details of its engagement by Bpa, as well as the stoichiometry and heterogeneity of the resulting complex, incompletely understood.

Multiple high-resolution structures of apo Bpa have been determined by X-ray crystallography and single-particle electron cryomicroscopy (cryo-EM)^25,26,29^. Bpa subunits adopt a four-helix bundle that further assembles into a 228 kDa homododecameric ring with 12-fold symmetry (Fig. 1a). In addition to the canonical dodecameric architecture, a previously determined crystal structure of a truncated Bpa variant (residues 44-153; PDB: 5IEU) revealed a tetrameric assembly stabilized by a distinct set of inter-subunit interactions, highlighting the intrinsic propensity of Bpa to adopt alternative oligomeric states under specific conditions.^25^ Consistent with this structural plasticity, we recently showed that full-length Bpa reversibly assembles into a dodecameric ring from dimeric and tetrameric species in a temperature-dependent manner^30^, underscoring the dynamic nature of its oligomeric equilibrium in solution. In that same study, we employed an unnatural surrogate substrate to circumvent the poor solubility of the native substrate (i.e. HspR) and identified a binding ratio of 12 Bpa subunits to 3 hTRF1 molecules. While these studies provided important insight into Bpa-mediated substrate recognition, reliance on an unnatural substrate leaves open the fundamental question of how Bpa engages its physiological targets. In particular, the stoichiometry and heterogeneity of the Bpa-HspR complex and the residues involved in substrate engagement remain incompletely understood.

**Fig 1.**
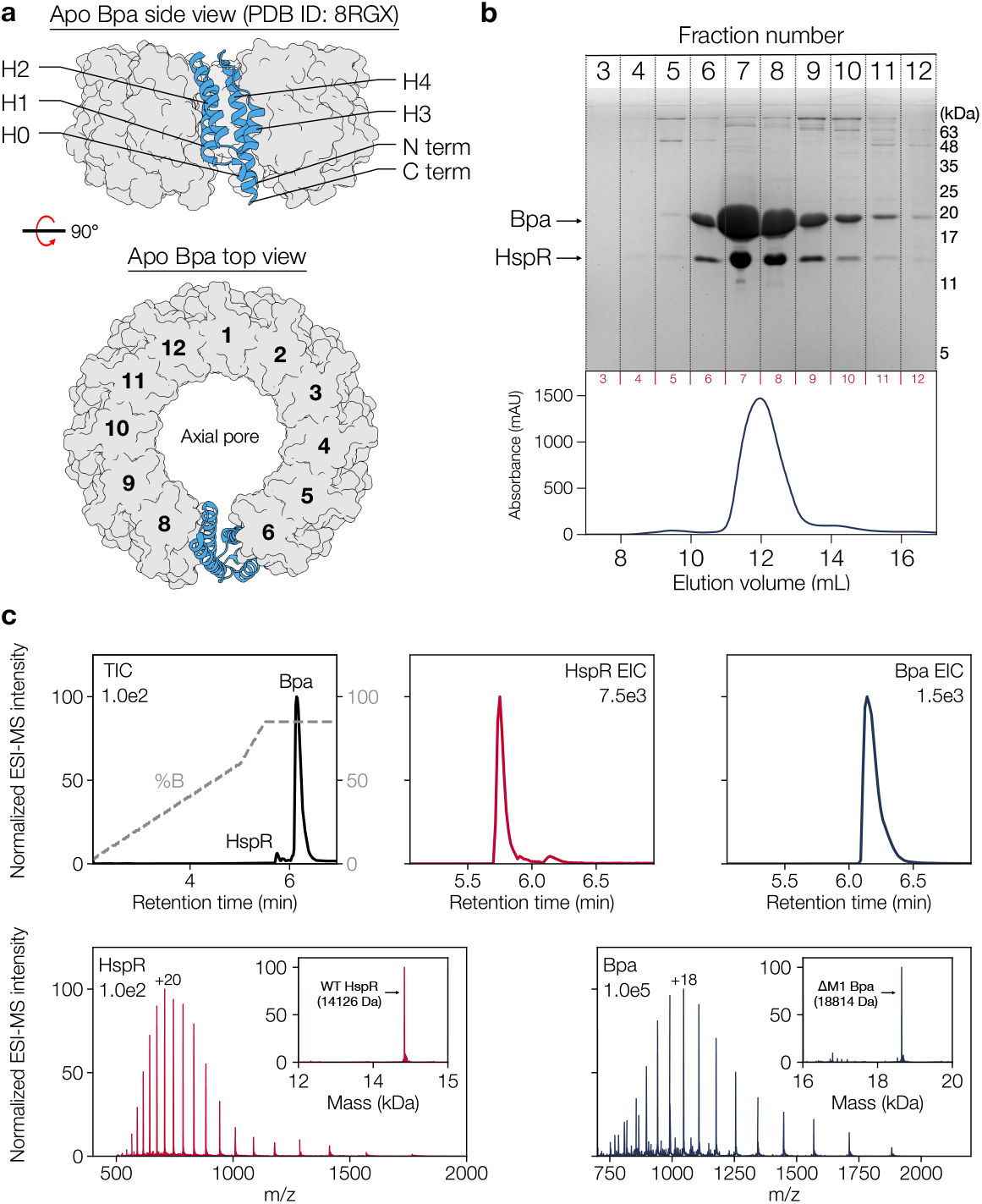
Structure of apo Bpa and native substrate HspR. (**a**) Side and top view of apo dodecameric Bpa (surface) highlighting one individual subunit within the ring (cartoon). PDB ID: 8RGX from van Rosen *et al*. 2025^29^. (**b**) SDS-PAGE and corresponding size exclusion chromatography of co-purified Bpa and HspR using a Superdex 200 Increase 10/300 GL column. Fraction numbers are indicated above each image. (**c**) Denaturing intact MS verifies the molecular weights of Bpa and HspR. The total ion chromatogram is overlaid with fraction of solvent B (i.e. acetonitrile) over time during reverse-phase liquid chromatography (dashed grey trace). The deconvoluted masses are within 1 Da of the theoretical mass for each protein^4^. Maximum ion intensities for each chromatogram or spectra are denoted.

Here, we use a combination of CDMS and cryo-EM coupled with biochemical analysis to characterize the interactions between Bpa and its native substrate, HspR.

Using CDMS and cryo-EM, we demonstrate the presence of both undecameric and dodecameric apo Bpa in solution and that the oligomeric form is impacted by the presence of a high-affinity substrate. CDMS shows a homogenous population corresponding to two HspR substrates per dodecameric Bpa, revealing a previously unknown complex stoichiometry. Finally, we were able to obtain a high-resolution map of a Bpa:HspR complex without the chemical crosslinkers used in previous structural studies^29^, revealing a defined substrate density localized to the C-terminal H4 helix of Bpa. Together, these results grant new insights into Bpa-mediated proteasomal degradation that inform the development of novel therapeutic strategies against *Mtb*.

## Results

### Protein purification and functional assays

We sought to reconstitute a stable Bpa:HspR complex suitable for structural characterization. A major challenge in assembling the complex is the insoluble and aggregation-prone nature of HspR, precluding its purification in a homogeneous, monodisperse form. In contrast, Bpa is well behaved in isolation and acts as the physiological binding partner for HspR. We reasoned that complex formation between Bpa and HspR could stabilize HspR in solution. We heterologously co-expressed untagged wild-type *Mtb* Bpa and His-SUMO-tagged *Mtb* HspR and used the His-tag to pull down the Bpa-HspR complex, followed by SEC (Fig. 1b). LC-MS analysis of the purified complex confirmed that Bpa and HspR are the only protein components present (Fig. 1c). The discrepancy in ion intensity between Bpa and HspR is due to their distinct ionization efficiencies, a well-known phenomenon in ESI^31^. Using a discontinuous protease assay, we showed that HspR is degraded in the presence of WT Bpa and WT 20S CP (Fig. S1). Conversely, a Bpa variant containing a Y173A substitution within the GQYL motif, which co-purified with HspR, failed to facilitate the degradation of HspR (Fig. S2). Similarly, the catalytically inactive 20S_βT1A_, when combined with WT Bpa, did not degrade HspR (Fig. S3). These data demonstrate that HspR degradation requires both Bpa and a functional 20S CP. Together, these results validate our co-expression strategy, confirm specific Bpa-dependent engagement of HspR, and provide a well-defined Bpa:HspR complex for structural characterization.

We aimed to obtain homogeneous apo Bpa to enable faithful comparisons between apo Bpa and the Bpa-HspR complex. Accordingly, both wild-type (WT) Bpa and Bpa_Δ155-166_ were purified under denaturing conditions to minimize the engagement of endogenous substrates from the expression host. The resulting purifications yielded highly homogenous Bpa suitable for structural characterization. To confirm proper function upon refolding, we performed a discontinuous protease assay. Refolded apo Bpa facilitated degradation of the unnatural substrate hTRF1 when combined with WT 20S CP (Fig. S4). No evidence of substrate degradation was observed when combined with catalytically inactive 20S_βT1A_ (Fig. S5).

### CDMS reveals co-existing stoichiometries of apo Bpa

We used CDMS to assess the stoichiometry of apo Bpa. CDMS determines the mass of individual ions by independently measuring their mass-to-charge ratio (*m/z*) and charge (*z*) as they oscillate within an electrostatic linear ion trap (ELIT)^10–12^. We employed a Waters Xevo™ CDMS instrument to analyze apo Bpa with an ion trapping time of 100 ms. We incubated apo Bpa at 37 °C overnight to promote self-assembly prior to CDMS analysis^30^. For apo Bpa, individual ions were plotted as a function of their measured *m/z* and *z* values. One-dimensional distributions along each axis were computed using kernel density estimation (KDE), which converts discrete single-ion measurements into smooth density profiles (see Methods). The resulting 2D *m/z* versus *z* plots revealed two distinct series of charge state distributions that nearly perfectly overlapped in the *m/z* dimension (Fig. S6) but are clearly separated by charge (Fig. 2a). As a result, these species cannot be resolved on the basis of *m/z* alone and are only distinguishable through direct charge measurement. This charge-based separation enabled unambiguous isolation of co-existing protein populations and the generation of accurate mass distributions for apo Bpa (Fig. 2c). Strikingly, the apo Bpa mass spectrum (blue trace) exhibited two well-defined peaks of comparable intensity at 209.8 and 229.3 kDa, corresponding to undecameric and dodecameric Bpa assemblies, respectively. Bpa is widely reported to assemble exclusively into a dodecamer and our previous native MS data acquired on a Q-TOF instrument relying solely on *m/z* information could only detect the dodecameric species with confidence^30^. Because undecameric and dodecameric Bpa differ by <10% in molecular weight and have nearly identical hydrodynamic radii and light-scattering properties, these species are not resolved by SEC-based or spectroscopic methods. Therefore, the emergence of the undecameric Bpa assembly highlights a previously unresolved stoichiometry that is uniquely revealed by direct charge measurement using CDMS.

**Fig 2.**
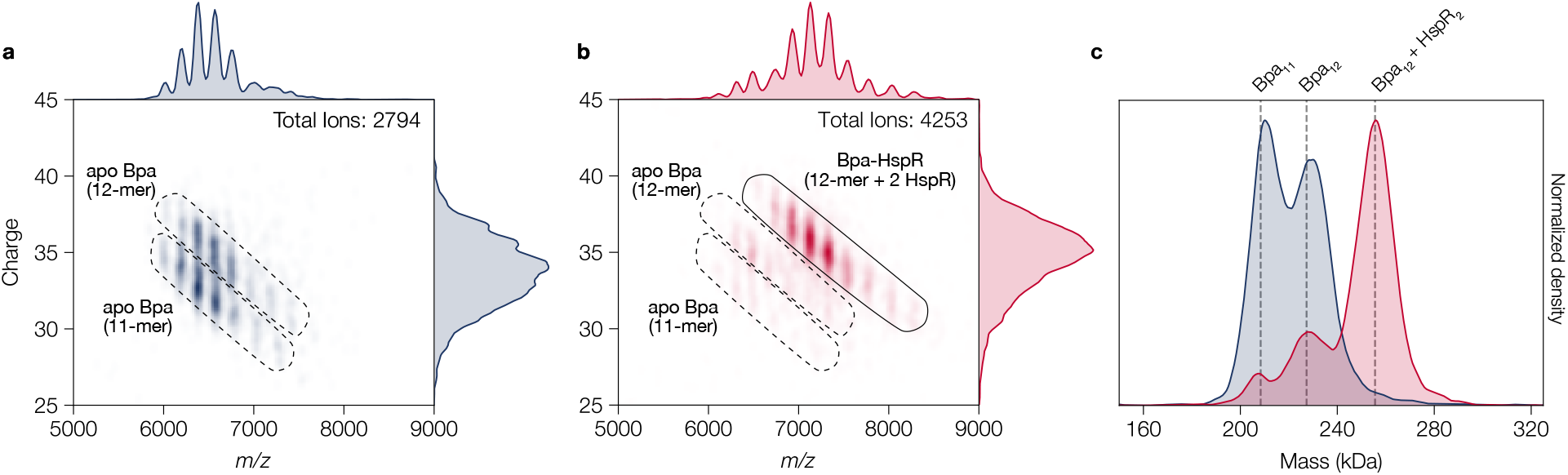
CDMS analysis of Bpa. 2D *m/z* versus *z* CDMS plots for (**a**) apo Bpa, and (**b**) Bpa-HspR complex (100 ms trapping time). Black solid and dashed lines indicate distinct populations displayed in the overlayed mass distributions (panel c). Bandwidth values for kernel density estimation of *z* and *m/z* were 0.5 and 60, respectively; (**c**) Mass distributions comparing apo and substrate bound Bpa:HspR data. The bandwidth value for kernel density estimation of mass was 5 kDa. Grey dashed lines represent the theoretical MW of each Bpa species. Each mass distribution was independently normalized to the highest density within each dataset.

### CDMS defines the stoichiometry of the Bpa-HspR complex

We repeated the CDMS measurements for the Bpa-HspR complex isolated by the pull-down strategy described above. The resulting 2D *m/z* versus *z* plots revealed that the two charge state distributions corresponding to apo undecameric and dodecameric Bpa, which dominate in the apo state, were markedly reduced but not fully depleted (Fig. 2b – dashed line), giving way to a dominant charge state distribution centred at higher *m/z* and *z* values (Fig. 2b – solid line). The signals corresponding to apo Bpa also nearly fully overlap with HspR-bound Bpa in the *m/z* space. As a result, these species cannot be resolved on the basis of *m/z* alone and are only distinguishable through direct charge measurement. Charge distributions show that the Bpa:HspR complex carries a higher average charge than apo Bpa (35.4 versus. 34.4) (Fig. 2a, b – right region of 2D plot). The corresponding mass distribution showed that the molecular species centred at 228.4 kDa was 27.4 kDa heavier relative to dodecameric apo Bpa at 255.8 kDa acquired within the same spectrum. This mass increase corresponds to 1.94 HpsR molecules bound to dodecameric apo Bpa, based on an HspR mass of 14.1 kDa (Fig. 2c). This gives rise to a Bpa_12_:HspR_2_ binding stoichiometry. Notably, we did not identify distinct mass population corresponding to an undecameric Bpa:HspR complex (e.g., Bpa_11_:HspR_2_, ~237 kDa; or Bpa_11_:HspR_3_, ~251 kDa), indicating that HspR preferentially associates with fully assembled dodecameric Bpa. All individual peaks in both the apo Bpa and Bpa:HspR mass distributions were well described by single Gaussian fits, though variation in peak widths (FWHM) suggests some underlying heterogeneity within each population (Fig. S7). We observed low-abundance shouldering in the Bpa:HspR profiles, possibly indicating minor heterogeneity in the number of substrates bound.

We previously showed that apo Bpa dissociates into lower-order species upon incubation at 4 °C^30^. Consistent with this, CDMS measurements revealed that apo Bpa incubated at 4 °C predominantly populates monomers, dimers, and tetramers (Fig. S8a, black trace). In contrast, HspR binding stabilizes higher-order Bpa oligomers even at low temperature (Fig. S8b, red trace), indicating that substrate engagement shifts the equilibrium toward assembled Bpa states independent of temperature (Fig. S8c). SEC-MALS analysis also showed stabilization of dodecameric Bpa upon HspR binding without incubation at physiological temperature (Fig. S9a, b). Although a shift in oligomeric equilibrium upon HspR binding was observed, SEC-MALS was unable to accurately determine the molecular weight of the complex due to its similar hydrodynamic radius and light-scattering properties to those of the apo dodecamer. This further highlights the capability of CDMS to produce unambiguous mass determinations for particles difficult to resolve using conventional methodologies such as SEC-MALS.

To further validate the stoichiometry of the Bpa:HspR complex, we next induced dissociation by heating the ion sampling inlet of a prototype charge detection mass spectrometer equipped with a similar ELIT analyzer but a distinct ion sampling interface from the commercial instrument used above. As a control, we first analyzed HspR-bound Bpa with the inlet at room temperature and an ion trapping time of 500 ms. Under these conditions, we measured ions with an average charge of 32.5 and an average mass of 256.7 kDa, consistent with an intact Bpa_12_:HspR_2_ complex (Fig. 3a). Increasing the inlet temperature to 300 °C resulted in a near complete dissociation of the complex, producing a distinct ion population with an average charge of 18.8 and an average mass of 224.6 kDa, corresponding to apo dodecameric Bpa. Notably, dodecameric Bpa retains only 58% of the total charge despite accounting for 87.5% of the Bpa_12_:HspR_2_ complex mass. The substantial charge reduction between the Bpa:HspR precursor and the apo Bpa product reflects asymmetric charge partitioning following HspR ejection upon ion activation (Fig. 3b)^32^. Due to the low molecular weight of the ejected HspR and the 500 ms ion trapping time, we did not detect signal corresponding to dissociated HspR in our CDMS experiments.

**Fig 3.**
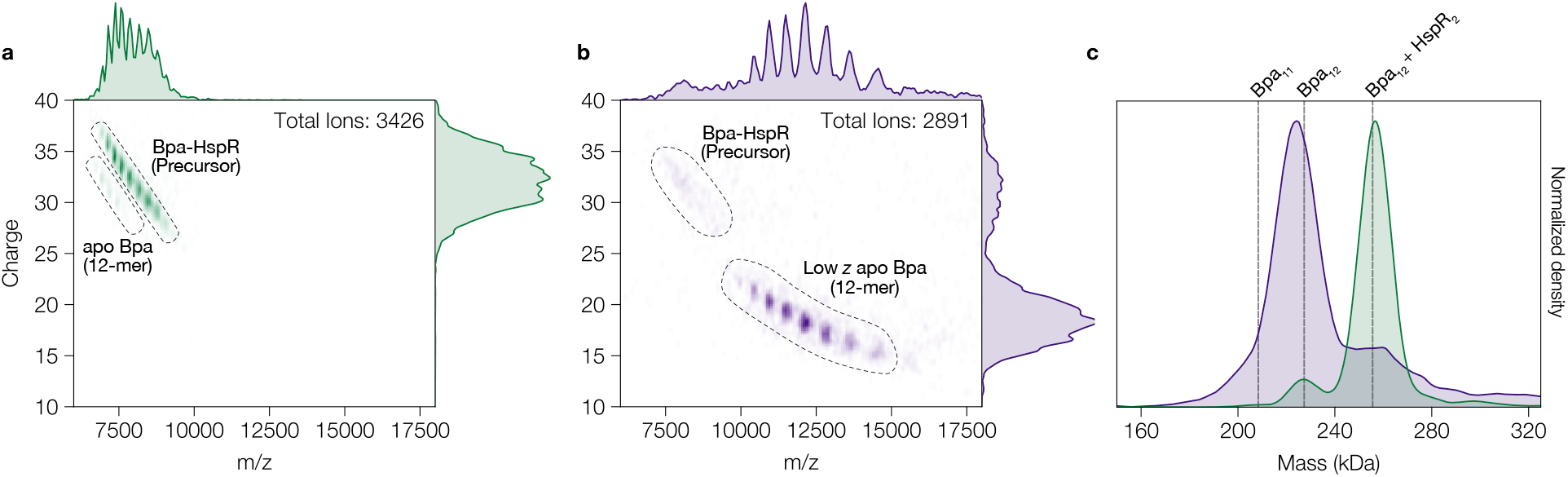
CDMS analysis of the Bpa-HspR complex with inlet heating (300 °C). Two-dimensional *m/z* versus *z* CDMS plots for Bpa-HspR measured with the ion sampling interface at (**a**) room temperature or (**b**) heated to 300 °C to induce ion activation. Black dashed lines indicate distinct populations displayed in the overlayed mass distributions (panel c). Bandwidth values for kernel density estimation of *z* and *m/z* were 0.5 and 60, respectively; (**c**) Mass distributions comparing gentle (room temperature) and activating (300 °C) conditions for Bpa:HspR data. The bandwidth value for kernel density estimation of mass was 5 kDa. Grey dashed lines represent the theoretical MW of each Bpa species. Each mass distribution was independently normalized to the highest density within each dataset.

Together, these CDMS measurements define the stoichiometry of the Bpa:HspR complex and demonstrate that HspR binding stabilizes higher-order Bpa assemblies independent of temperature. Critically, the near-complete overlap of the corresponding *m/z* spectra underscores that these conclusions would be inaccessible to conventional native MS and spectroscopic approaches (Fig. S9ab) and highlights the unique power of CDMS to resolve complex, heterogeneous assemblies through direct, single-particle charge measurement.

### Cryo-EM reveals undecameric apo Bpa and maps Bpa–HspR interaction sites

We used single-particle cryo-EM to reconstruct three-dimensional (3D) maps of apo Bpa_Δ155-166_ and the Bpa_Δ155-166_:HspR complex in order to examine the structural basis of substrate engagement. The particles used for this analysis were derived from an experiment in which Bpa_Δ155-166_ or Bpa_Δ155-166_:HspR were incubated with the 20S CP. Bpa_Δ155-166_ exhibits increased affinity for the 20S CP and was used to increase fraction bound^25^. Following particle classification, a large population of unbound Bpa (apo and Bpa:HspR) was identified and used for reconstructions displayed here. Our initial cryo-EM analysis of apo Bpa predominantly recovered fully assembled dodecameric rings, along with several lower-resolution classes for which the oligomeric state could not be determined conclusively. However, given the CDMS measurements indicated the presence of an undecameric Bpa population, we modified our *ab-initio* reconstruction parameters to account for higher resolution details and improve the recovery of a smaller undecameric particle population. Subsequent nonuniform refinement in C1 symmetry yielded reconstructed maps for apo undecameric Bpa (4.7 Å), apo dodecameric Bpa (4.0 Å), and the native Bpa:HspR complex (3.9 Å) (Fig. 4a-c). In all three maps, the structured four-helix bundle comprising H1, H2, H3, and the upper region of H4 spanning residues S111 to approximately F138, is resolved at modest to high local resolution (Figs. S10 and S11). In contrast, local resolution decreases toward the dynamic C terminus of H4 in all three maps. Although the maps are not sufficient for detailed atomic modeling in this region, they clearly differentiate the number of subunits incorporated into the Bpa ring structure (Fig. 4a, b). Classifications on the Bpa:HspR dataset did not recover an undecameric Bpa:HspR class, consistent with the CDMS data indicating that HspR binding favors the dodecameric assembly. Additional unassigned density is observed near the C-terminal region of H4 in both undecameric and dodecameric maps (Fig. S12). One possible explanation for this density is interactions between adjacent H4 helices within the ring structure, previously shown to promote tetramer formation in the absence of substrate and under cold incubation conditions^25,30^. Importantly, apo Bpa was purified under denaturing conditions and refolded on-column prior to analysis, minimizing the likelihood of co-purified contaminants. The observed density is localized specifically between adjacent H4 helices in a pattern consistent across both undecameric and dodecameric maps, rather than distributed randomly, as would be expected for heterogeneous contaminants.

**Figure 4.**
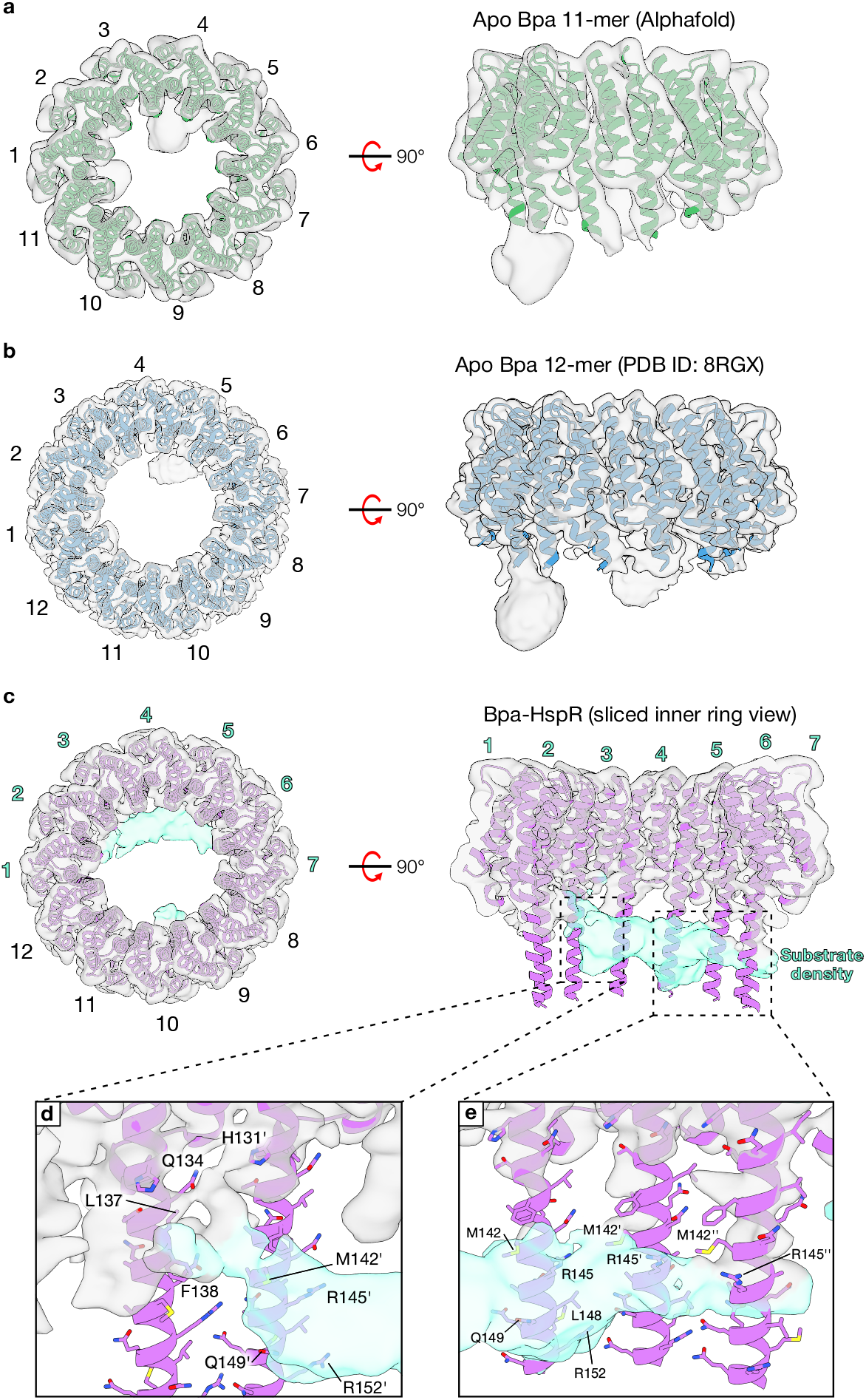
Cryo-EM analysis of apo undecameric Bpa, apo dodecameric Bpa, and Bpa:HspR. Top (left) and side view (right) of the cryo-EM maps for (**a**) apo undecameric Bpa (4.7 Å) and (**b**) apo dodecameric Bpa (4.0 Å). An AlphaFold3 predicted structure of Bpa_Δ155-166_ and experimentally determined apo dodecameric Bpa structure (PDB ID: 8RGX) have been rigid-body fit into the undecameric and dodecameric maps, respectively. The number of subunits is indicated. (**c**) Top (left) and inner ring view (right) of the density map for Bpa:HspR (3.9 Å). An AlphaFold3 predicted structure of Bpa_Δ155-166_ has been rigid-body fit, with residues predicted to be disordered hidden. The density map is coloured based on proximity to the rigid-body fit model, with teal representing unassigned density corresponding to bound substrate. Subunits denoted with teal labels in the top and inner ring views highlight interactions between Bpa and HspR, which are shown in greater detail in panels (**d**) and (**e**). Residues participating in substrate engagement are labelled.

The co-expression construct coupled with our pull-down strategy yielded a stable Bpa:HspR complex with high HspR occupancy in the absence of cross-linking reagents, enabling visualization of substrate-associated density under near-native conditions. Following non-uniform refinement in C1 symmetry, particles were subjected to focused 3D classification using a mask over the C-terminal region of Bpa to improve visualization of the bound HspR density (Fig. S11e). Two classes displaying consistent extra density were combined to generate a final map with clear substrate-associated density in the lower lumen of the Bpa ring (Fig. 4c teal). This substrate-associated density in the Bpa:HspR complex is distinct from the density observed in the apo Bpa and attributed to the possible H4-H4 helices interactions, as supported by analogous focussed 3D classification performed on dodecameric apo Bpa (Fig. S10).

As the C-terminal regions of Bpa is intrinsically flexible, available experimental structures of dodecameric Bpa are not resolved beyond residue L147^25,26,29^. To aid the interpretation of the density extending into this poorly resolved region, we fit an AlphaFold3 model of dodecameric Bpa into the cryo-EM map. For regions resolved in our cryo-EM structures, the AlphaFold3 model agrees well with crystallographic Bpa models (Fig. S13). The AlphaFold3 model enabled examination of Bpa:HspR contacts beyond regions resolved in current experimental structures. The substrate-associated density spans the lower lumen of Bpa and is positioned adjacent to two groups of subunits within the dodecameric ring, corresponding to subunits 2–3 and 4–6 (Fig. 4c teal labels). At the 2-3 interface, this density is localized near residues L137 and F138 (Fig. 4d), with weaker density extending upward toward Q134 and H131’, suggestive of additional interactions. Additional density is observed further along helix H4, in the vicinity of residues M142’, R145’, Q149’, and R152’, consistent with substrate engagement extending into the flexible C-terminal region and beyond the resolved range of available structures. Following the lumen-spanning substrate density, a similar interaction pattern is observed at the 4-6 interface (Fig. 4e). While density near M142 and R145 is present across all subunits, a more pronounced density is observed proximal to the C terminus of subunit 5, with residues L148, Q149, and R152 embedded within the surrounding density and thus likely contributing to HspR engagement. Together, these data localize HspR binding to the lower lumen of Bpa and support a contribution of the C-terminal H4 region to substrate engagement.

## Discussion

Previous studies have demonstrated that Bpa-dependent degradation of HspR is essential for *Mtb* virulence^19,20,20,28^. However, the exact interactions that mediate complex formation, as well as complex stoichiometry have remained elusive. Although prior investigations have established a strong understanding of apo Bpa and its interactions with substrates, most have relied on heavily truncated constructs of both Bpa and HspR to enable characterization^22,24,25,29^. Although detailed insights into Bpa substrate engagement have been obtained using an unnatural model substrate (hTRF1)^30^, whether these conclusions extend to the native substrate HspR remained unclear. This work aimed to examine Bpa and HspR interactions under near native conditions without the use of cross-linking reagents. Here, we utilized a construct that co-expressed tagged HspR and untagged Bpa to co-purify the intact complex. This co-purification strategy allowed us to overcome issues arising from the aggregation-prone HspR and produce stable, native complexes suitable for characterization. Using CDMS, we identified a previously unreported complex stoichiometry of Bpa_12_:HspR_2_ using full-length Bpa and HspR. By contrast, we observed a Bpa_12_:hTRF1_3_ complex stoichiometry using NMR experiments^30^. The difference in size between HspR (14.1 kDa) and the DNA-binding domain of hTRF1 (6.6 kDa) may account for substrate stoichiometry differences within the Bpa ring structure. Interestingly, the presence of HspR appears to shift the oligomeric equilibrium toward fully assembled dodecamers independent of incubation temperature^30^.

Following CDMS-based structural characterization, we used single-particle cryo-EM to examine apo Bpa_Δ155-166_ and Bpa_Δ155-166_:HspR complexes and obtained density maps for both apo undecameric and dodecameric Bpa_Δ155-166_. The observation of an undecameric species further underscores the oligomeric plasticity of Bpa, which is known to exist as dimeric, tetrameric, and dodecameric species in solution^24,25,27,29,30^. Our analysis also yielded the highest-resolution density map of a Bpa:HspR complex to date. The density corresponding to bound HspR spans the lower lumen of Bpa, contacting multiple subunits. However, all sites of interaction appear localized to the C-terminal region of H4, spanning residues 130-155. This region contains multiple hydrophobic (L137, F138, M142, L148) and charged (R145, R152) residues forming bands within the lower internal ring structure of Bpa. Consistent with these observations, von Rosen et al.^29^, examining a cross-linked Bpa_36-139_:HspR_ΔC9_ complex by cryo-EM, identified H131, F138, and R145 as key HspR-interacting residues^29^. In addition, residues I133, L137, and M142 exhibited chemical shift perturbations upon binding to the unnatural substrate hTRF1 in NMR experiments^30^. Taken together, these data corroborate previously identified interaction sites (H131, F138, R145) under near-native conditions and without chemical crosslinking, while additionally revealing contributions from residues further along the C-terminal region of H4 (L148, Q149, R152) that extend beyond the range resolved in existing dodecameric structures.

The use of two complementary single-particle detection techniques offered this work a unique synergy and enabled a detailed structural characterization. The ability of CDMS to unambiguously resolve highly similar oligomeric species within a seemingly homogeneous sample provided invaluable insights for cryo-EM, which relies heavily on particle classification strategies. The identification of low-abundance oligomeric states via CDMS enabled the recovery of an unprecedented undecameric Bpa species that would have been otherwise masked by the dodecameric species. Our cryo-EM maps also validate that the unusual stoichiometries observed in CDMS data do occur in solution and are not artifacts of electrospray ionization, ion transmission, or ion activation. Together, these highly complementary methodologies provide a deeper understanding of Bpa and its interactions with native substrate HspR. Targeting regions responsible for substrate engagement could serve as an interesting strategy to disrupt the essential *Mtb* proteasome system. Furthermore, the absence of a eukaryotic Bpa homolog only increases its attractiveness as a potential target for a novel therapeutic^22,25,27^.

## Acknowledgements

B.T.V.D. acknowledges support from a Graduate Tuition Scholarship from the University of Guelph. A.B.U acknowledges support from the Fonds de Recherche du Québec-Santé (FRQ-S) doctoral fellowship. Financial support was provided by Canadian Institutes of Health Research Project Grant PJT451412 to S.V., as well as PJT419240 and PJT195698 to M.M, by a Discovery Grant from the Natural sciences and Engineering Council of Canada (RGPIN/03031-2022) to N.Z, (RGPIN-2021-02843) to S.V, and (RGPIN-2018-06070) to M.M, and by the Princess Margaret Cancer Foundation (PMCF) to M.M. The Centre de Recherche en Biologie Structurale (CRBS) is funded by Fonds de Recherche du Québec (Health Sector) Research Centres Grant #288558. We thank K. Basu, and K. Sears at the Facility for Electron Microscopy Research (FEMR) at the McGill University for their assistance with microscope operation and data collection. FEMR is supported by the Canadian Foundation for Innovation, Quebec Government, and McGill University. The denaturing MS data were recorded at the Mass Spectrometry Facility of the Advanced Analysis Centre, University of Guelph. We thank Dr. Dyanne Brewer (University of Guelph) for assistance with MS measurements. We thank Dr. Algirdas Velyvis (University of Guelph) for guidance and helpful discussions.

## Statement of contribution

B.T.V.D. and S.V. initiated the project; B.T.V.D., A.B.U., A.H., A.F.A.K., M.M., K.G., N.Z., and S.V. designed and performed research, analyzed data, and wrote the paper; B.T.V.D., A.H., J.U., D.B., K.R., K.G., and S.V. contributed new reagents/analytic tools.

## Data availability

Electron microscopy density maps have been deposited in the Electron Microscopy Databank (accession nos. EMD-76916, EMD-76918, EMD-76919). CDMS data have been deposited to the MassIVE database as entry MSV000101635.

