## Supplementary Information for "Resolving hidden stoichiometries in Bacterial proteasome activator (Bpa)-substrate complexes by cryo-EM and charge detection mass spectrometry"

This file contains:

**Materials and Methods**
**Figure S1.** WT Bpa-HspR, WT 20S CP discontinuous degradation assay
**Figure S2.** Y173A Bpa-HspR, WT 20S CP discontinuous degradation assay
**Figure S3**. WT Bpa-HspR, T1A 20S CP discontinuous degradation assay
**Figure S4.** Refolded WT Bpa, WT 20S, hTRF1 discontinuous degradation assay
**Figure S5.** Refolded WT Bpa, T1A 20S, hTRF1 discontinuous degradation assay
**Figure S6.** Overlapping apo Bpa 11-mer and 12-mer populations during CDMS
**Figure S7.** Peak fitting of apo Bpa and Bpa:HspR mass distributions
**Figure S8.** Temperature-independent dodecameric assembly due to HspR binding
**Figure S9.** SEC-MALS analysis of assemble apo Bpa and Bpa:HspR
**Figure S10.** Single-particle cryo-EM data refinement process for apo Bpa
**Figure S11.** Single-particle cryo-EM data refinement process for Bpa:HspR
**Figure S12.** C-terminal density indicating adjacent Bpa subunit interactions
**Figure S13.** Structural alignment of Bpa models with an Alphafold3 predicted structure

#### Materials and Methods

*Plasmids and constructs*

Codon-optimized genes encoding for Bpa (UniProt: *bpa*, P9WKX3), HspR (UniProt: *hspr,* O06302), proteasome α-subunit (UniProt: *prcA*, P9WHU1), and proteasome β-subunit excluding the propeptide (UniProt: *prcB*, P9WHT9) were synthesized and inserted into kanamycin-resistant pET24-based vectors (Biobasic, Markham, ON, Canada). The β-subunit construct contains a C-terminal TEV-SUMO-$His_6$ tag, while all other constructs include a N-terminal $His_6$-SUMO affinity tag. The Bpa-HspR coexpression construct was produced using Gibson assembly[1]. The coexpression construct consists of HspR bearing an N-terminal $His_6$-SUMO and untagged Bpa. Point mutations for Bpa Y173A variants were introduced via Quickchange site directed mutagenesis [2].

*Protein expression and purification*

The Bpa:HspR coexpression constructs (WT, Δ155-166, Y173A) and 20S CP subunits were expressed and purified under non-denaturing conditions. Plasmids were transformed into chemically competent NEB T7 Express *LysY E. coli* cells. Cells were grown in lysogeny broth (LB) to a $OD_{600}$ 0.6-0.8 before induction with 0.2 mM Isopropyl β-d-1-thiogalactopyranoside (IPTG). All 20S CP subunits were expressed overnight at 16 °C while shaking at 180 rpm. Bpa:HspR was expressed for 3 hours at 37 °C shaking at 180 rpm before cell harvesting. For the purification of all proteins, cells were harvested via centrifugation at 5,000 × *g* for 20 minutes. The cell pellets were frozen for future purification. All cells were resuspended in 30 mL of lysis buffer and lysed via sonication (Heat Systems INC Sonicator XL 2020 Ultrasonic Liquid Processor) using a 5-minute program consisting of 10 seconds on and 20 seconds off. IMAC buffers for Bpa-HspR consisted of 50 mM Tris, 600 mM KCl, supplemented with either 20 (lysis/wash), 50 (strong wash), or 500 (elution) mM imidazole at pH 7.0. The lysis for Bpa:HspR also contained 10 mM adenosine-5'-triphosphate disodium salt and 10 mM $MgCl_2$ to prevent the endogenously expressed ATP-dependent chaperone DnaK from binding HspR. The 20S CP IMAC buffers consisted of 50 mM Tris, 300 mM NaCl, supplemented with either 20 (lysis/wash), 50 (strong wash), or 500 (elution) mM imidazole at pH 7.0. Endpoints for the strong wash and elution steps were determined via Bradford assay. Eluted proteins were dialyzed overnight in lysis buffer + 1mM dithiothreitol (DTT) with the appropriate proteases to cleave the affinity tags. 20S CP was assembled using the protocol described by Turner *et al.* (2025).[3] After dialysis and cleavage, a second IMAC step was conducted to remove cleaved affinity tags. Size exclusion chromatography (SEC) using either a Superose 6 or Superdex 200 (S200) was performed using an ÄKTA PURE™ chromatography system as a final purification step. The final protein products for Bpa-HspR were spiked with 10% glycerol (v/v) and flash frozen in liquid $N_2$ before storage at -80 °C. 20S CP constructs were never frozen and stored at 4 °C until further use.

Multiple Bpa constructs (WT, Δ155-166, Y173A) used for CDMS and single-particle cryo-EM analysis were transformed, grown, expressed (induced with 0.2 mM IPTG and expressed overnight at 16 °C while shaking at 180 rpm), and harvested as described above. The purification was then performed under denaturing conditions to ensure minimal endogenous substrate binding within apo Bpa samples. Cells were resuspended and lysed in 30 mL of denaturing lysis buffer consisting of 6M Gdn-HCl, 50 mM HEPES and 20 mM imidazole, pH 7.0, as described above. Initial IMAC was performed under denaturing conditions consisting of two washes using lysis buffer and 2 strong wash steps using lysis buffer containing 50 mM imidazole. Before eluting, the protein was refolded on column via the addition of buffer containing 50 mM HEPES, 600 mM KCl and 20 mM imidazole. To ensure minimal concentrations of residual Gdn-HCl the column was washed with 200 mL of refolding buffer. Once refolded and washed, endpoints for the strong wash (50 mM imidazole) and elution (500 mM imidazole) steps were determined via Bradford assay. All refolded Bpa constructs were never frozen and stored at 4 °C until further use. Refolded Bpa displayed the same elution volumes during SEC as Bpa purified under non-denaturing conditions indicating proper refolding. Activity of refolded Bpa was also tested using degradation assays which resulted in Bpa-depenedent degradation of non-native substrate (Fig S4 and S5).

Protein quality for all stages of IMAC and SEC were assessed using sodium dodecyl sulfate polyacrylamide gel electrophoresis (SDS-PAGE). Final concentrations of all protein stocks were determined spectrophotometrically using a NanoDrop 2000 spectrophotometer. All extinction coefficients for concentration measurements were calculated via ExPASy's ProtParam (https://web.expasy.org/protparam/).

*Discontinuous SDS-PAGE-based protein degradation assays*

Degradation assays containing Bpa-HspR (WT or Y173A) and 20S CP (WT or βT1A) were performed at 37 °C for 48 hours. Protein concentration for these reactions consisted of 20 μM Bpa-HspR (Bpa monomer) and 2 μM 20S CP (α-β pair**)**. Degradation assays containing refolded Bpa (WT), 20S CP (WT, T1A), and hTRF1 were performed at 37 °C for 48 hours. Protein concentration for these reactions consisted of 10 μM Bpa (Bpa monomer), 2 μM 20S CP (α-β pair), and 10 μM hTRF1. Aliquots were taken at discrete time points and the reaction was quenched by mixing sample with Laemlli buffer (4:1) followed by subsequent heat inactivation at ~95 °C for 5 minutes[4]. Visualization of protein degradation was performed using SDS-PAGE. 20 μL of sample was loaded and run into 20% acrylamide gels for all samples to ensure adequate visualization. BlueElf Prestained Protein Marker (5-245kDa). Imaging of all gels was conducted on a BioRad ChemiDoc Imaging System (Hercules, CA).

*SEC-MALS*

WT Bpa and Bpa:HspR was subjected to SEC-MALS analysis using an OMNISECmulti-detector SEC system (Malvern Panalytical, United Kingdom) fitted with OMNISEC RESOLVE and OMNISEC REVEAL modules. 100 μL samples were injected at 2mg/mL and loaded onto a P3000 Protein SEC column (300 × 8mm, Malvern Panalytical) equilibrated in 150 mM NaCl, 50 mM Tris-HCl, pH 7.5 at a flow rate of 1 mL/min. Molecular weight was calculated using light scattering detectors at 90° (right-angle light scattering) and 7° (low-angle light scattering). BSA was used as a standard for molecular weight calibration.

*Charge Detection Mass Spectrometry*

50 μL aliquots of apo Bpa and Bpa:HspR were individually buffer exchanged into 100 mM ammonium acetate (Invitrogen, AM9070G) using Rad Bio-Spin® P-6 size-exclusion columns (Bio-Rad, 7326221). Protein concentrations of the buffer-exchanged proteins samples were then determined using a ThermoScientific NanoDrop One (Thermo

Scientific, 840274100) prior to sample preparation. Aliquots of the buffer-exchanged apo Bpa were incubated at either 4 or 37 °C overnight prior to measurement. Bpa:HspR was thawed, buffer exchanged and run within the same day. All samples were diluted to a final Bpa monomer concentration of 1 μM using 100 mM ammonium acetate before subsequent measurements. Mass analysis was performed using a Waters Xevo™ CDMS instrument with an electrostatic linear ion trap (ELIT). The heated inlet data was collected on a prototype instrument in which the ion sampling interface could be heated to >300 °C. For both the prototype and Xevo, the ELIT device is based on the design by the Jarrold group at Indiana University, which has been described previously[5]. 10 μL of sample was loaded into a reusable glass emitter (5 μm internal diameter), and ions were generated by positive-ion-mode static nanoelectrospray ionization (nanoESI). Ions were trapped in the ELIT for 100 ms and the frequency and amplitude information were converted to m/z and z values, respectively, and ultimately mass (m/z × z). The glass emitter was cleaned extensively between injections using LC-MS grade water and biological cleaning solution to prevent sample carryover. Individual ion measurements were recorded as discrete (*m/z*, *z*) data points and visualized as two-dimensional scatter plots. One-dimensional distributions of mass, *m/z*, and *z* were computed using kernel density estimation (KDE), which produces smooth density profiles from discrete single-ion data. KDE bandwidth parameters are specified in each figure legend. Data were collected until a minimum of approximately 3000 ions were recorded in total. Signal processing and data visualization were performed using in-house Python 3.15.5 scripts.

*Denaturing Intact Mass Spectrometry*

Reverse phase liquid chromatography (LC) on a BEH™ C4 (1.7 µM, 2.1 ×150 mm) ACUITY™ column (Waters) connected to a Waters I-class ultra-performance liquid chromatography (UPLC™) System was used to separate 10 pmol of Bpa:HspR (Bpa monomer). At a flow rate of 0.2 mL/min, proteins were separated using a 13-minute H2O: acetonitrile (ACN) gradient with a column temperature of 40 °C. Both mobile phases were acidified using 0.1% [v/v] formic acid. The ACN content was increased linearly from 2–85 % over 7 minutes, prior to a single sawtooth gradient where ACN was ramped from 2–85 % over 3 minutes before a final flush of 2 % ACN to equilibrate for the next injection. The LC eluent was directed to a SYNAPT™ G2-Si quadrupole time-of-flight (Q-TOF) mass spectrometer fit with a standard dual emitter electrospray ionization (ESI) source. Measurements were conducted in positive-ion mode with the capillary voltage set to +3 kV. The TOF mass analyzer was set to resolution mode, and surveyed precursor ions from 50–2,000 Th using 0.4 second scans. External calibration was performed with the "ESI-L low concentration tune mix" (G1969-8500–Agilent Technologies, CA). The instrument was also dynamically calibrated via infusing 2 µM Leucine Enkephalin (peptide sequence YGGFL; 1+ m/z 556.2771) dissolved in 50% [v/v] ACN and 0.1% [v/v] formic acid at 10 µL/min from the lock mass sprayer, which was sampled every 20 seconds. Mass spectral analysis and deconvolution were performed using MassLynx™ v4.2 Software and the MaxEnt1 function, respectively. Mass deconvolutions were reanalyzed and confirmed using UniDec 8.1.2[6].

*Electron cryomicroscopy sample preparation and data collection*

For sample preparation, complexes were reconstituted with $20S_{T1A}$ core particle in the presence of $Bpa_{\Delta155\text{-}166}$ or $Bpa_{\Delta155\text{-}166}$-HspR for the two datasets. They were reconstituted in a 1:3 ratio (8 µM $20S_{T1A}$ + 24 µM $Bpa_{\Delta155\text{-}166}$ or $Bpa_{\Delta155\text{-}166}$-HspR (Bpa dodecamer)). Each sample was applied to a holey carbon grid (C-flat CF-2/1-3Cu-T) that had been glow discharged in air at 5 mA for 15 s. Sample vitrification was performed in a Vitrobot Mark IV (Thermo Fisher Scientific) at 4 °C and 100% humidity. Each sample was incubated on the grid for 3 s and the grid was blotted for 4 s with a blot force of 1 before plunging in liquid ethane.

Datasets were collected using SerialEM software[1] in the Titan Krios at FEMR-McGill. Movies were recorded on a Gatan K3 direct electron detector equipped with a Quantum LS imaging filter. The total dose used for each movie was 50 e/Å$^2$, equally spread over 30 frames. The datasets were collected at a magnification of 105,000x, yielding images with a calibrated pixel size of 0.855 Å. The nominal defocus range used during data collection was between -1.25 µm and -2.75 µm. Data collection parameters for all the datasets are summarized in Table S1.

*Electron cryomicroscopy data processing*

All cryo-EM data processing was performed using cryoSPARCv4[2]. Movies were corrected for beam-induced motion correction using patch motion correction, and the CTF parameters were estimated using patch CTF estimation. Micrographs with CTF fit resolution that was worse than 10 Å, and those with unusually high full-frame motion were discarded at this stage. The remaining micrographs were used for particle picking using blob picker with a minimum and maximum diameter of 50 Å and 150 Å respectively to select the apo-Bpa/Bpa-HspR particles. These particles were subjected to two rounds of 2D classification, and the classes resembling apo-Bpa/Bpa-HspR were used as templates for template picker. The template-based picked particles were subjected to two rounds of 2D classification for particle curation. The particles from the best classes were then used for ab-initio reconstruction and hetero-refinement. The class(es) displaying the characteristic structural features of the Bpa-ring were selected, and subjected to reference-based motion correction, global and local CTF refinement[3] in cryoSPARC, and then used for high-resolution non-uniform refinement. These particles were further classified using focussed 3D classification with a mask over the C-terminal H4 helices of Bpa to probe for HspR-binding. In case of the Bpa-HspR dataset, classes that showed strong extra density were pooled and subjected to non-uniform refinement. Local resolution was estimated for all the maps using cryoSPARCv4. ChimeraX[4] was used for visualizing the maps and figure preparation. Data processing details for all the datasets are summarized in Supplementary figures 10 to 11.

#### Supplementary figures

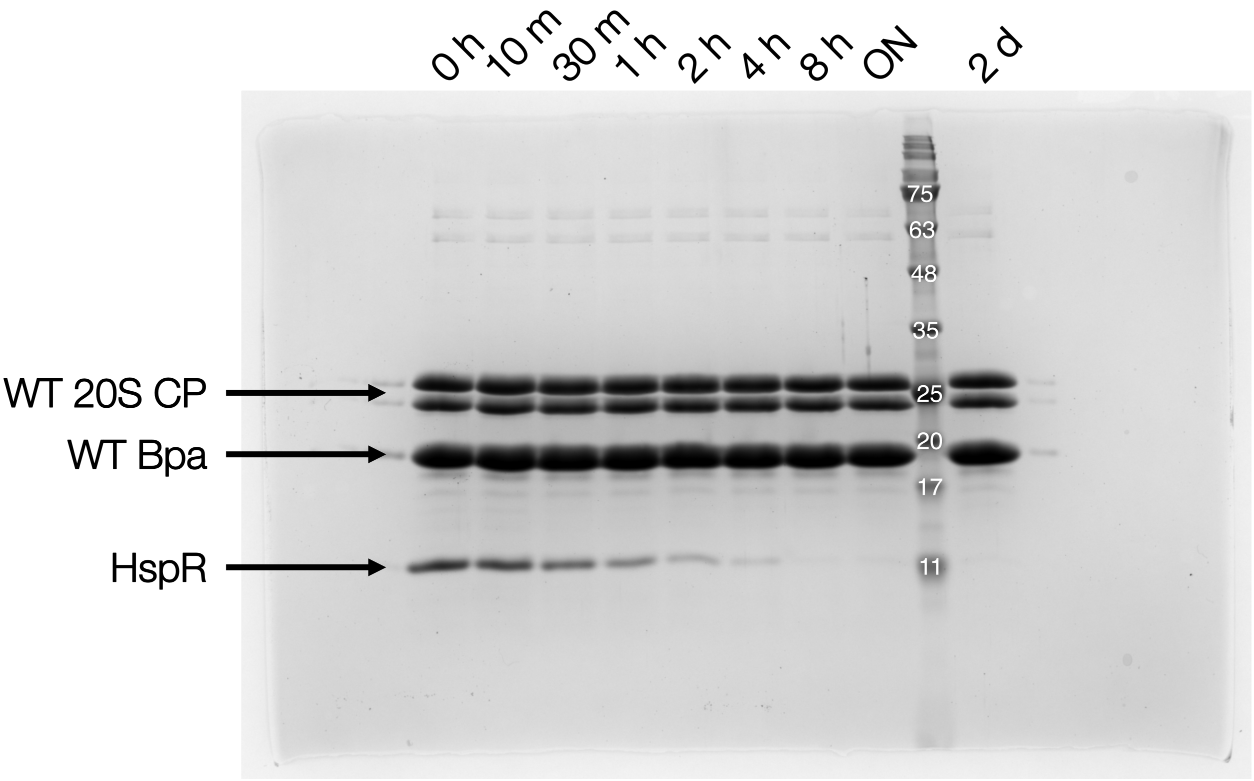

**Fig S1. Bpa-mediated proteasomal degradation of native substrate HspR.** All components (WT Bpa-HspR complex, WT 20S CP) were combined and incubated at 37 °C for 48 hours. Aliquots were taken at discrete time points (0 h – 48 h) and quenched via dilution of sample into Laemlli buffer (4:1) before subsequent heat inactivation at ~95 °C for 5 minutes. Reaction protein concentrations were 2 μM 20S CP (α-β pair) and 20 uM Bpa-HspR (Bpa monomer). 20% acrylamide gels were used for SDS-PAGE analysis.

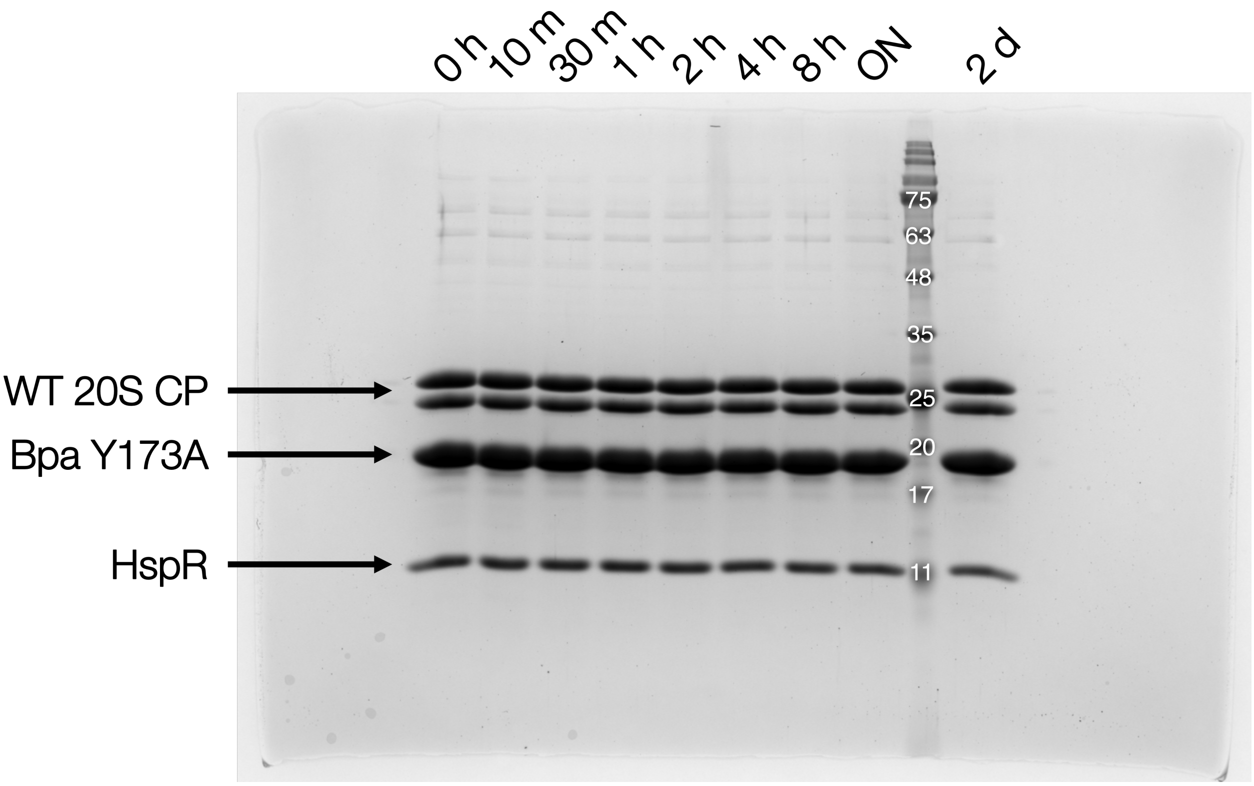

**Fig S2. Bpa Y173A abrogates association with 20S CP resulting in inhibited HspR degradation.** Lack of degradation with GQYL deficient Bpa indicates Bpa-mediated degradation of HspR. All components (Y173A Bpa-HspR complex, WT 20S CP) were combined and incubated at 37 °C for 48 hours. Aliquots were taken at discrete time points (0 h – 48 h) and quenched via dilution of sample into Laemlli buffer (4:1) before subsequent heat inactivation at ~95 °C for 5 minutes. Reaction protein concentrations were 2 μM 20S CP (α-β pair) and 20 uM Bpa-HspR (Bpa monomer). 20% acrylamide gels were used for SDS-PAGE analysis.

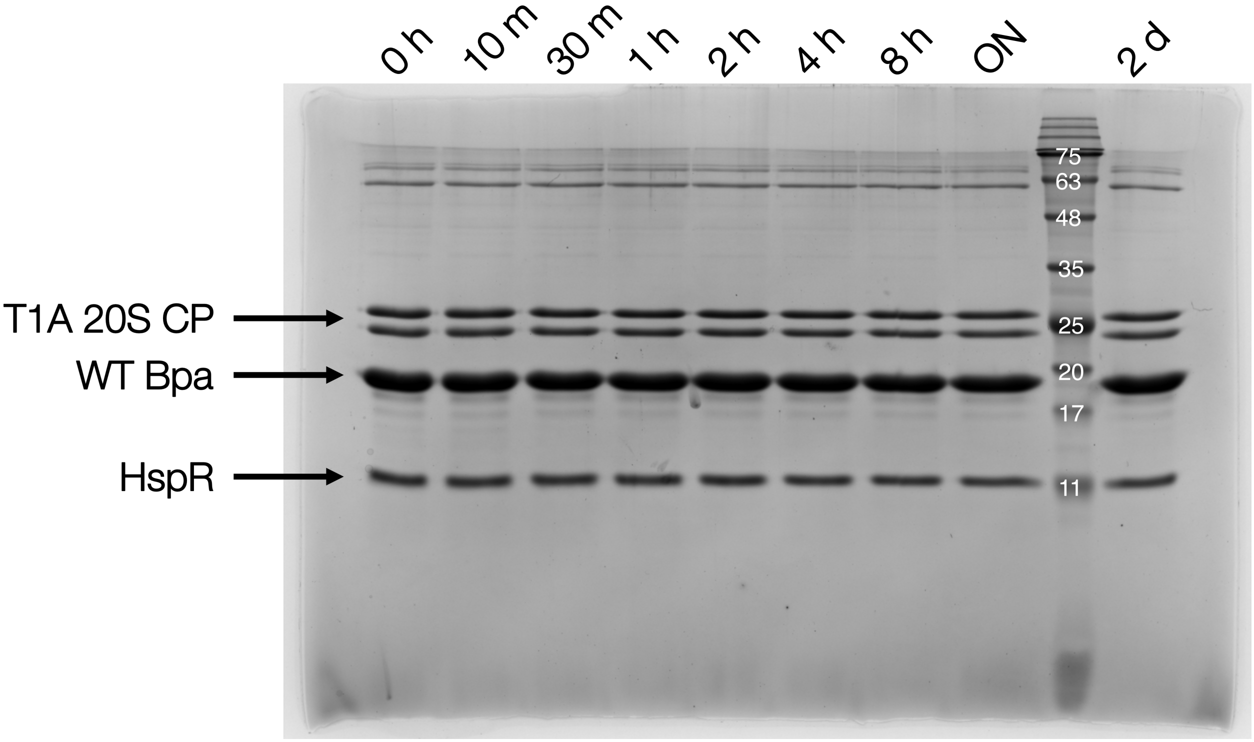

**Fig S3. No degradation of HspR occurs with catalytically inactive 20S CP.** All components (WT Bpa-HspR complex, T1A 20S CP) were combined and incubated at 37 °C for 48 hours. Aliquots were taken at discrete time points (0 h – 48 h) and quenched via dilution of sample into Laemlli buffer (4:1) before subsequent heat inactivation at ~95 °C for 5 minutes. Reaction protein concentrations were 2 μM 20S CP (α-β pair) and 20 uM Bpa-HspR (Bpa monomer). 20% acrylamide gels were used for SDS-PAGE analysis.

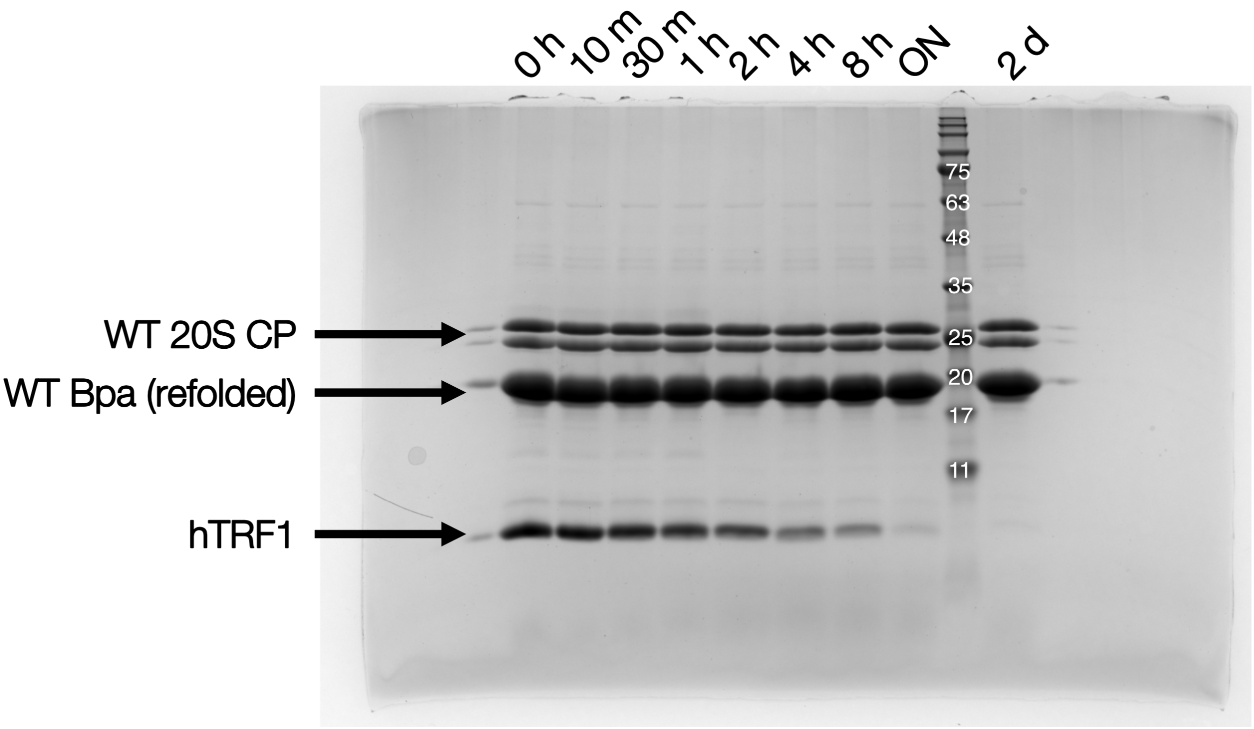

**Fig S4. Refolded WT Bpa is able to facilitate degradation of hTRF1.** All components (refolded WT Bpa, WT 20S CP, hTRF1) were combined and incubated at 37 °C for 48 hours. Aliquots were taken at discrete time points (0 h – 48 h) and quenched via dilution of sample into Laemlli buffer (4:1) before subsequent heat inactivation at ~95 °C for 5 minutes. Reaction protein concentrations were 2 μM 20S CP (α-β pair) and 10 μM (Bpa monomer), and 10 μM hTRF1. 20% acrylamide gels were used for SDS-PAGE analysis.

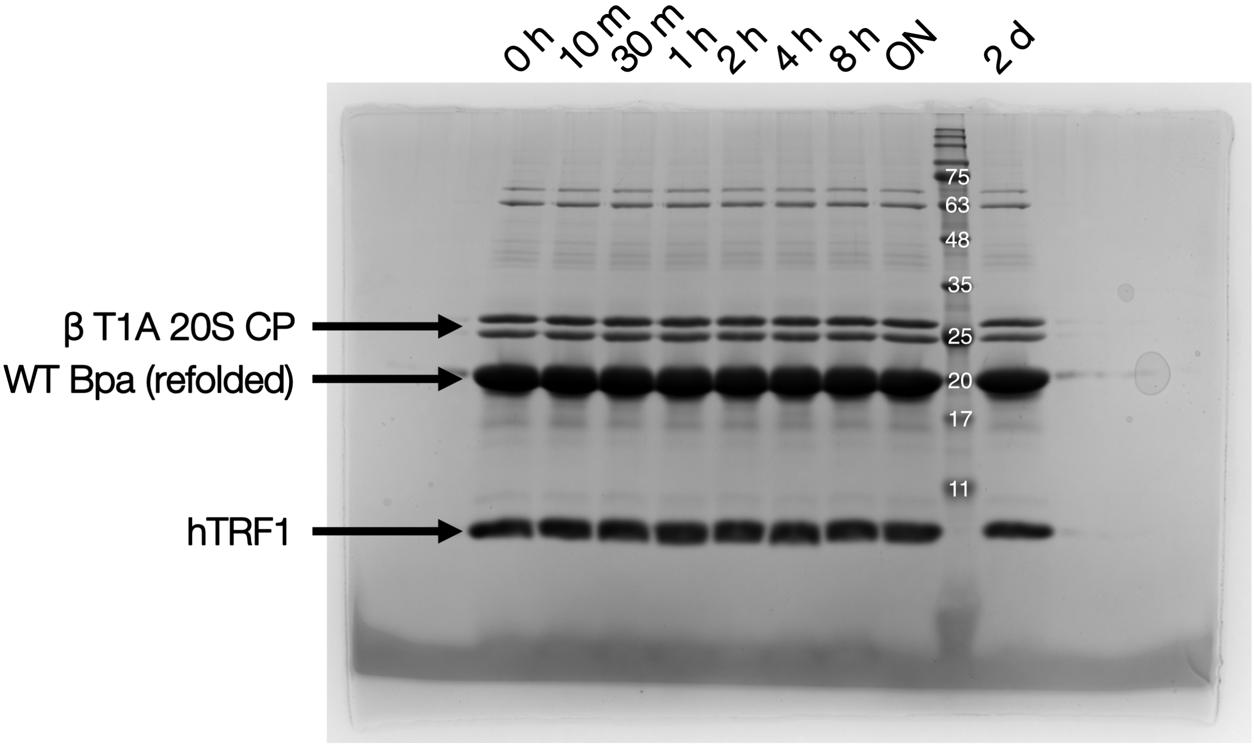

**Fig S5. No degradation of hTRF1 when refoled WT Bpa is combined with inactive 20S CP.** All components (refolded WT Bpa, T1A 20S CP, hTRF1) were combined and incubated at 37 °C for 48 hours. Aliquots were taken at discrete time points (0 h – 48 h) and quenched via dilution of sample into Laemlli buffer (4:1) before subsequent heat inactivation at ~95 °C for 5 minutes. Reaction protein concentrations were 2 μM 20S CP (α-β pair) and 10 μM (Bpa monomer), and 10 μM hTRF1. 20% acrylamide gels were used for SDS-PAGE analysis.

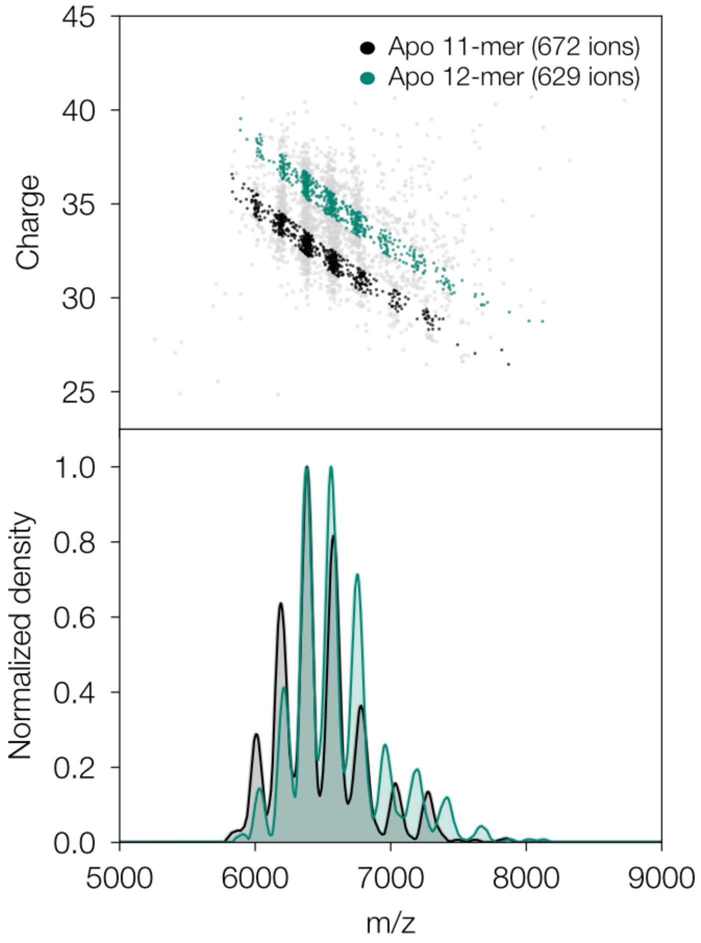

**Fig S6. Apo Bpa 11-mer and 12-mer populations present nearly identical *m/z* distributions.** Upper panel displays the selected populations for undecameric (black) and dodecameric (green) Bpa. To generate the population-specific *m/z* distributions, individual ions were assigned to the undecameric or dodecameric population based on their measured mass: ions with masses within ±2% of the mean undecameric (209.8 kDa) or dodecameric (229.3 kDa) peak mass were selected for each respective trace. The lower panel displays a comparative *m/z* distribution of both the 11-mer and 12-mer. The bandwidth value for kernel density estimation of *m/z* was 60. Each *m/z* distribution was independently normalized to the highest density within each dataset.

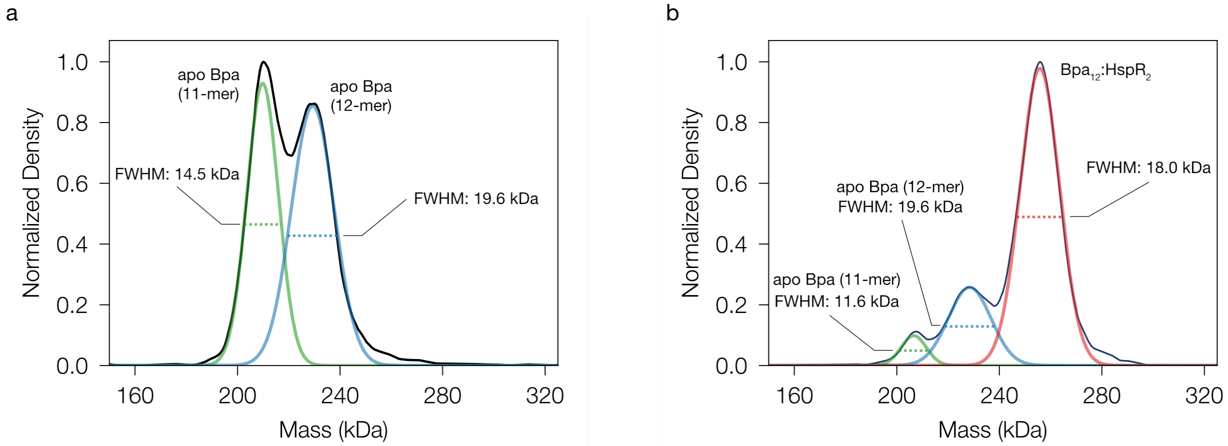

**Fig S7. Peak fitting of Bpa mass distribution reveals homogenous populations**. KDE distributions of mass for (a) apo Bpa and (b) Bpa:HspR. The bandwidth value for kernel density estimation of mass was 5 kDa. Each population within the respective sample was fit using a single Gaussian function. The populations and corresponding FWHM values for each population are denoted. All experiments were conducted at 1 μM Bpa (monomer) in 100 mM ammonium acetate, pH 7.0.

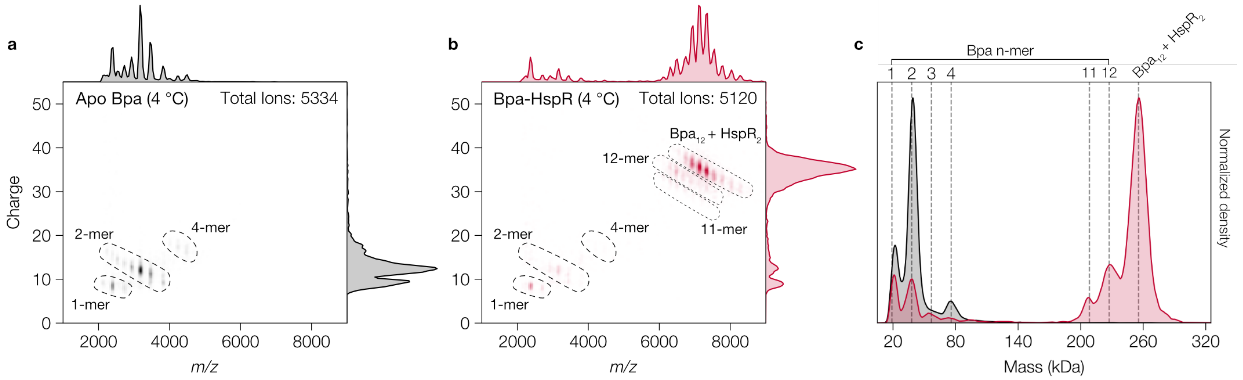

**Fig S8. HspR aids in Bpa assembly when compared to unassembled Bpa (4 °C).** Bpa:HspR was not incubated at 37 °C to promote assembly yet remains predominantly in larger oligomers due to the presence of HspR**.** Two-dimensional *m/z* vs *z* CDMS plots for (**a**) unassembled apo Bpa, and (**b**) Bpa:HspR complex (100 ms trapping time). Black dashed lines are labelled and indicate distinct populations displayed in the overlayed mass distributions (panel c). Bandwidth values for kernel density estimation of *z* and *m/z* were 0.5 and 60, respectively; (**c**) Mass distributions comparing apo and Bpa:HspR data. The bandwidth value for kernel density estimation of mass was 5 kDa. Grey dashed lines represent the theoretical MW of each Bpa species. Each mass distribution was independently normalized to the highest intensity within each dataset.

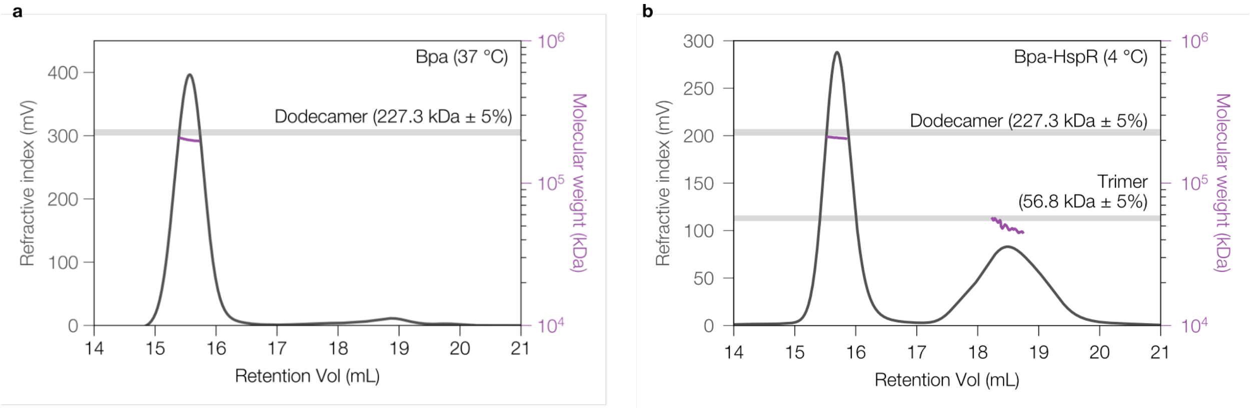

**Fig S9. SEC-MALS analysis shows no difference molecular weight between apo and bound Bpa.** Chromatograms and corresponding molecular weights for (**a**) apo Bpa (37 °C) and (**b**) Bpa:HspR (4 °C). Apo Bpa was incubated at 37 °C overnight to promote assembly. Bpa-HspR was stored at 4 °C prior to analysis. Both samples were then analyzed at 20 °C. A single predominant peak corresponding to dodecameric Bpa is observed. The left y-axis and grey trace represent the refractive index signal. The right y-axis and purple trace indicate the estimated molecular weight. Grey bars denote ±5% of the expected molecular weight for each indicated oligomeric species. An OMNISEC multi-detector SEC system (Malvern Panalytical, United Kingdom) fitted with an OMNISEC RESOLVE and OMNISEC REVEAL modules was used.

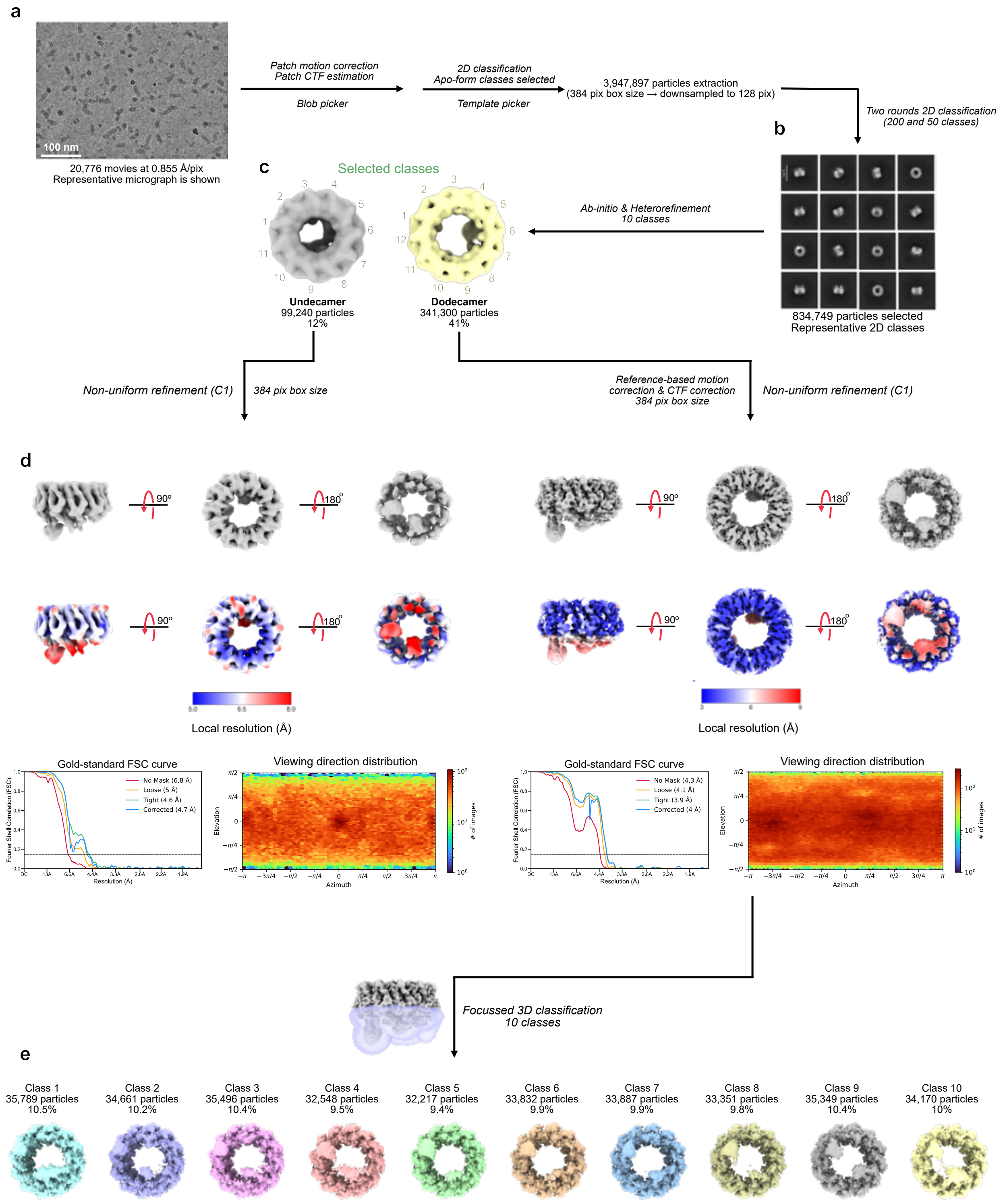

**Fig S10. Single-particle cryo-EM data refinement process for apo undecameric and dodecameric Bpa.** (a) Representative micrograph. (b) Representative 2D classifications. (c) 3D classes after heterogenous refinement for undecameric (left) and dodecameric (right) Bpa. (d) Final consensus maps after non-uniform C1 refinement for undecameric (left) and dodecameric (right) Bpa with Gold-Standard Fourier shell correlation plots under each respective sample. (e) Focused 3D refinement on the C-terminal region of Bpa resulting in 10 classes (Mask used for focussed refinement is shown in violet as a transparent surface).

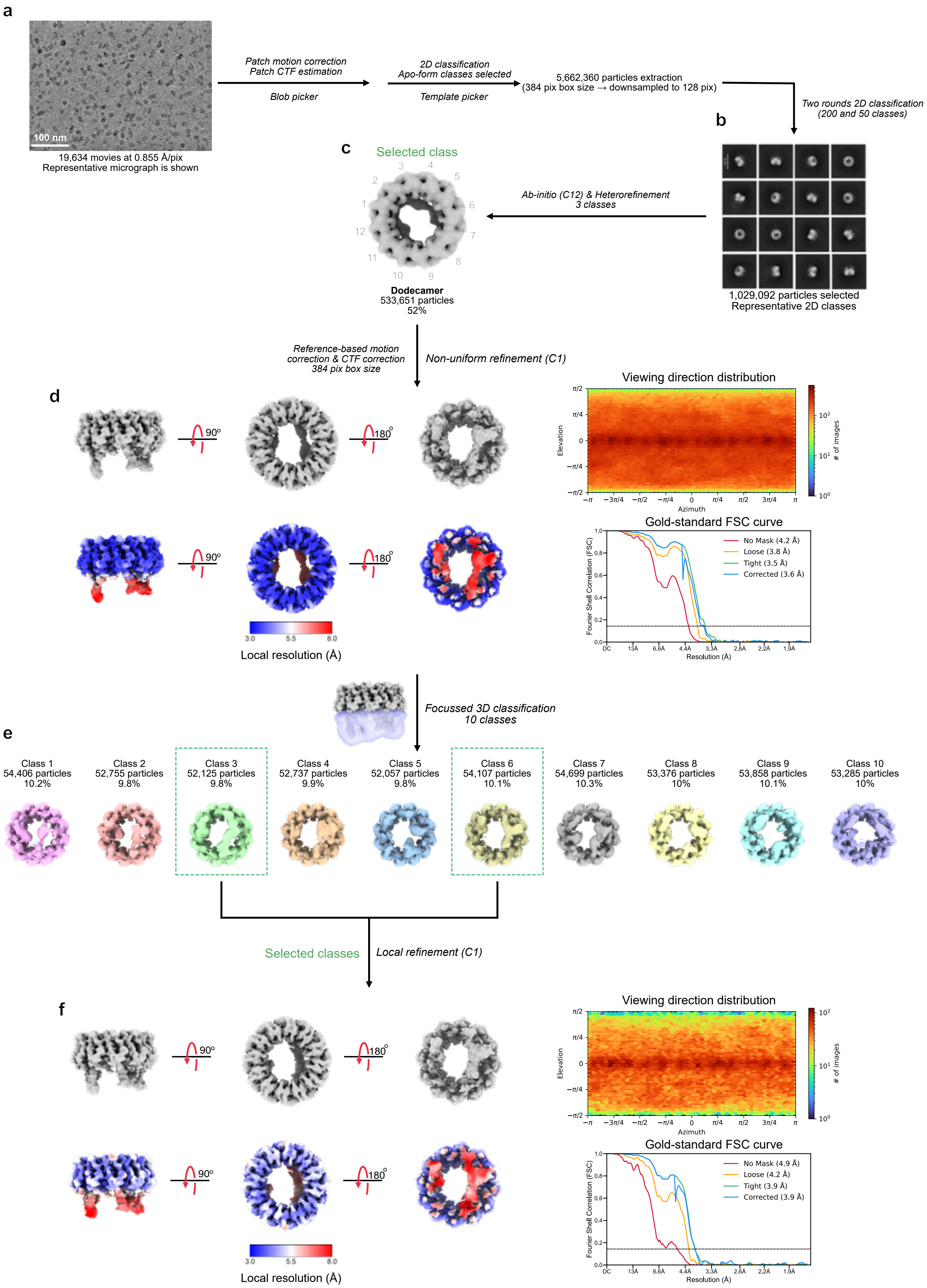
a
100 nm
19,634 movies at 0.855 Å/pix
Representative micrograph is shown
Patch motion correction
Patch CTF estimation
Blob picker
2D classification
Apo-form classes selected
Template picker
5,662,360 particles extraction
(384 pix box size → downsampled to 128 pix)
Two rounds 2D classification
(200 and 50 classes)
b
1,029,092 particles selected
Representative 2D classes
Ab-initio (C12) & Heterorefinement
3 classes
c
Selected class
Dodecamer
533,651 particles
52%
Reference-based motion
correction & CTF correction
384 pix box size
Non-uniform refinement (C1)
d
90°
180°
Local resolution (Å)
Viewing direction distribution
Elevation
Azimuth
### of images
Gold-standard FSC curve
No Mask (4.2 Å)
Loose (3.8 Å)
Tight (3.5 Å)
Corrected (3.6 Å)
Fourier Shell Correlation (FSC)
Resolution (Å)
Focussed 3D classification
10 classes
e
Class 1
54,406 particles
10.2%
Class 2
52,755 particles
9.8%
Class 3
52,125 particles
9.8%
Class 4
52,737 particles
9.9%
Class 5
52,057 particles
9.8%
Class 6
54,107 particles
10.1%
Class 7
54,699 particles
10.3%
Class 8
53,376 particles
10%
Class 9
53,858 particles
10.1%
Class 10
53,285 particles
10%
Selected classes
Local refinement (C1)
f
90°
180°
Local resolution (Å)
Viewing direction distribution
Gold-standard FSC curve
No Mask (4.9 Å)
Loose (4.2 Å)
Tight (3.9 Å)
Corrected (3.9 Å)

**Fig S11. Single-particle cryo-EM data refinement process for Bpa:HspR.** (**a**) Representative micrograph. (**b**) Representative 2D classifications. (**c**) 3D classes after heterogenous refinement for Bpa:HspR. (**d**) Final consensus maps after non-uniform C1 refinement for Bpa:HspR with Gold-Standard Fourier shell correlation plots to the right. (**e**) Focused 3D refinement on the C-terminal region of Bpa:HspR resulting in 10 classes (Mask used for focussed refinement is shown in violet as a transparent surface). (**f**) Consensus map for class 3+6 after C1 refinement with Gold-Standard Fourier shell correlation plots to the right.

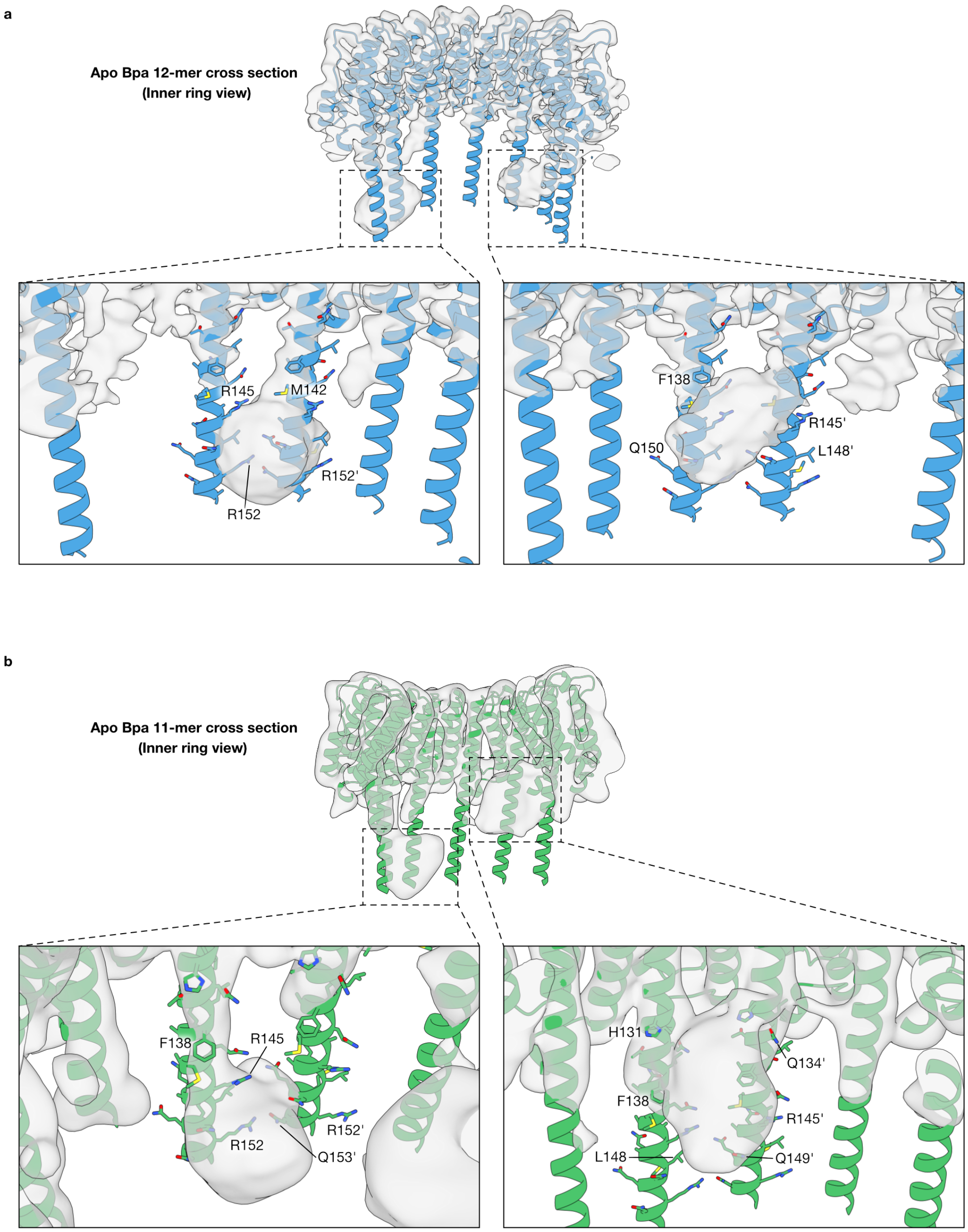

**Fig S12. Unassigned density in dynamic C-terminal regions of Bpa could indicate interactions between adjacent H4 helices.** Density maps for (**a**) dodecameric and (**b**) undecameric apo Bpa show clustered density between adjacent subunits. An experimentally determined cryo-EM structure (PDB: 8RGX) and an Alphafold3 predicted undecamer have been rigid body fit into the density map for dodecameric and undecameric Bpa, respectively. Zoomed regions for both oligomers highlight the clustered density in the lower H4 region. Image generation was conducted using ChimeraX 1.10[7].

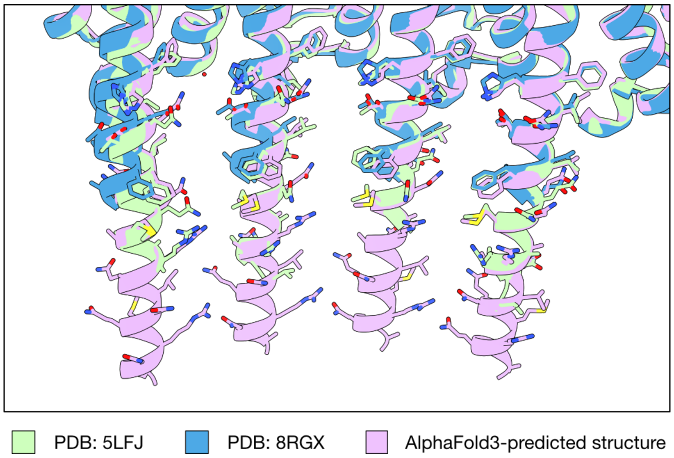

**Fig S13. Experimental dodecameric Bpa structures (PDB ID: 5LFJ, 8RGX) aligned to an AlphaFold3 predicted structure.** The RMSD for the alignment of 5LFJ and 8RGX to the predicted structure are 0.436 and 0.446 Å, respectively. Side chains for all models are displayed. Alignment and image generation were conducted using ChimeraX 1.10[7].

Table S1. Data collection and refinement statistics for cryo-EM.

| | **Apo Bpa (12-mer) Consensus map** | **Apo Bpa (11-mer) Consensus map** | **Bpa:HspR Class 6+3** |
|---|---|---|---|
| EMDB ID | EMD-76916 | EMD-76918 | EMD-76919 |
| **Data collection and processing** | | | |
| Microscope and camera | FEI Titan Krios with Gatan K3 Camera | FEI Titan Krios with Gatan K3 Camera | FEI Titan Krios with Gatan K3 Camera |
| Magnification (×) | 105,000 | 105,000 | 105,000 |
| Voltage (kV) | 300 | 300 | 300 |
| Data acquisition software | Serial EM | Serial EM | Serial EM |
| Electron dose (e-/$Å^2$) | 50 | 50 | 50 |
| Defocus range (um) | -1.75 to 2.75 | -1.75 to 2.75 | -1.75 to 2.75 |
| Pixel size (Å) | 0.855 | 0.855 | 0.855 |
| Number of micro-graphs | 20,776 | 20,776 | 19,634 |
| Processing soft-ware | cryoSPARC v4 | cryoSPARC v4 | cryoSPARC v4 |
| Symmetry imposed | C1 | C1 | C1 |
| Initial particle im-ages (no.) | 3,947,897 | 3,947,897 | 5,662,360 |
| Final particle im-ages (no.) | 341,300 | 99,240 | 106,232 |
| Map resolution (at FSC=0.143) (Å) | 4 | 4.7 | 3.9 |
